# Cold-induced [Ca^2+^]_cyt_ elevations function to support osmoregulation in marine diatoms

**DOI:** 10.1101/2022.03.11.483981

**Authors:** Friedrich H. Kleiner, Katherine E. Helliwell, Abdul Chrachri, Amanda Hopes, Hannah Parry-Wilson, Trupti Gaikwad, Nova Mieszkowska, Abdul Chrachri, Thomas Mock, Glen L. Wheeler, Colin Brownlee

## Abstract

Diatoms are a group of microalgae that are important primary producers in a range of open ocean, freshwater and intertidal environments. The latter can experience significant long- and short-term variability in temperature, from seasonal variations to rapid temperature shifts caused by tidal immersion and emersion. As temperature is a major determinant in the distribution of diatom species, their temperature sensory and response mechanisms likely have important roles in their ecological success. We have examined the mechanisms diatoms use to sense rapid changes in temperature, such as those experienced in the intertidal zone. We find that the diatoms *Phaeodactylum tricornutum* and *Thalassiosira pseudonana* exhibit a transient cytosolic Ca^2+^ ([Ca^2+^]_cyt_) elevation in response to rapid cooling, similar to those observed in plant and animal cells. However, [Ca^2+^]_cyt_ elevations were not observed in response to rapid warming. The kinetics and magnitude of cold-induced [Ca^2+^]_cyt_ elevations correlate with the rate of temperature decrease. We do not find a role for the [Ca^2+^]_cyt_ elevations in enhancing cold tolerance, but show that cold shock induces a Ca^2+^-dependent K^+^ efflux and reduces mortality of *P. tricornutum* during a simultaneous hypo-osmotic shock. As inter-tidal diatom species may routinely encounter simultaneous cold and hypo-osmotic shocks during tidal cycles, we propose that cold-induced Ca^2+^ signalling interacts with osmotic signalling pathways to aid in the regulation of cell volume. Our findings provide insight into the nature of temperature perception in diatoms and highlight that cross-talk between signalling pathways may play an important role in their cellular responses to multiple simultaneous stressors.

## Introduction

Diatoms are a group of silicified unicellular algae that represent one of the most important primary producers in modern oceans. They are abundant in diverse marine environments, most notably in polar and temperate upwelling regions, where they play a critical role at the base of the marine food web (Malviya et al., 2016). Diatom communities are abundant across a broad temperature range in the surface ocean from sea ice to tropical oceans. Diatoms are also important primary producers in freshwater and brackish ecosystems, where they likely encounter an even greater range of temperatures (Souffreau et al., 2010).

Global rises in surface temperature due to anthropogenic CO_2_ emissions are set to have profound influence on marine ecosystems (Gattuso et al., 2015). These future changes in our climate will also increase the variability of temperature regimes and the prevalence of extreme events, such as marine heatwaves, that may co-occur with other stressors such as low pH or deoxygenation (Harley et al., 2006; Smale et al., 2019; Gruber et al., 2021). Understanding the physiological response of diatoms and other marine phytoplankton to changes in global temperature regimes is therefore of the utmost importance. Temperature has an important impact on diatom cell physiology, influencing cell size and formation of the silica frustule (Montagnes and Franklin, 2001; Svensson et al., 2014; Javaheri et al., 2015). Individual species display a thermal niche with distinct temperature growth optima that reflect their natural environment (Liang et al., 2019). The upper and lower thermal tolerance limits, rather than the optima themselves, appear to have the greatest influence on the distribution of individual diatom species (Anderson and Rynearson, 2020), with temperatures in excess of the upper thermal tolerance limits leading to a rapid increase in the rates of cell death (Baker and Geider, 2021).

Many of these studies have focussed on the physiological responses of diatoms to longer term changes in temperature. However, diatoms will also experience short term temperature variations within their natural habitat. This is particularly so for those species that inhabit intertidal rocky shores or estuarine habitats where immersion and emersion is associated with rapid and regular temperature fluctuations. Rapid temperature changes are potentially highly damaging to diatom cells, demonstrated by their much greater vulnerability to abrupt rather than gradual temperature increases (Souffreau et al., 2010). Temperature variability may also have an important influence on the ability of diatoms to adapt to their thermal niche, as *Thalassiosira pseudonana* exhibited accelerated adaptation to higher temperatures under a fluctuating temperature regime (Schaum et al., 2018). Despite the importance of thermal tolerance in diatom physiology and ecology, relatively little is known about the physiological mechanisms that allow diatoms to perceive and respond to changes in temperature, particularly during short-term fluctuations.

Many of the cellular mechanisms involved in temperature sensing in eukaryotes involve temperature-induced changes in the structure of nucleic acids, proteins or biological membranes that lead to a range of downstream physiological responses (Sengupta and Garrity, 2013). Ca^2+^-dependent signalling mechanisms play an important role in these temperature sensing pathways. In animal cells, heat stress is associated with Ca^2+^ influx into the cytosol via the TRPV family of temperature-sensitive ion channels (Xu et al., 2002; Clapham and Miller, 2011). Ca^2+^ signalling also plays a role in sensing low temperature in animals, for example underpinning the rapid cold hardening response of insects (Teets et al., 2013). Land plants also employ Ca^2+^ signalling mechanisms in their response to both low and high temperatures. Rapid cooling of plants induces a transient cytosolic Ca^2+^ ([Ca^2+^]_cyt_) elevation, which leads to changes in gene expression and the establishment of cold tolerance (Knight et al., 1996; Tahtiharju et al., 1997; Knight and Knight, 2012). Some plants, such as the moss *Physcomitrium*, also display [Ca^2+^]_cyt_ elevations in response to heat shock (Saidi et al., 2009). In other plants, such as *Arabidopsis*, high temperatures do not induce [Ca^2+^]_cyt_ elevations, although Ca^2+^ elevations are observed within the chloroplast (Lenzoni and Knight, 2019). Potential temperature sensors in plants include the cold sensitive COLD1/RGA1 complex in *Oryza sativa*, which is proposed to either function as a Ca^2+^ channel or to activate other Ca^2+^ channels (Ma et al., 2015). Specific cyclic nucleotide-gated ion channels and annexins may also play a role in temperature sensing pathways, with mutant strains in *Physcomitrium, O. sativa* and *Arabidopsis* exhibiting diminished [Ca^2+^]_cyt_ elevations in response to cold- and heat shock (Cui et al., 2020; Liu et al., 2021). However, it is currently unclear whether these ion channels sense temperature directly or are activated indirectly, e.g. through changes in membrane rigidity (Plieth et al., 1999) or the cytoskeleton (Pokorna et al., 2004).

Our understanding of Ca^2+^ signalling in diatoms remains in its infancy, although Ca^2+^- dependent signalling mechanisms have been identified in response to a range of environmental stimuli, such as the supply of nutrients (phosphate and iron), hypo-osmotic shock and the detection of toxic aldehydes (Falciatore et al., 2000; Vardi et al., 2006; Helliwell et al., 2021; Helliwell et al., 2021). Initial experiments using *Phaeodactylum tricornutum* cells expressing the bioluminescent Ca^2+^ reporter aequorin did not detect [Ca^2+^]_cyt_ elevations in response to low (4 °C) or high (37 °C) temperature (Falciatore et al., 2000). More recently, genetically-encoded fluorescent Ca^2+^ reporters have been successfully expressed in *P. tricornutum* and *T. pseudonana*, enabling high resolution imaging of [Ca^2+^]_cyt_ elevations in single diatom cells (Helliwell et al., 2021; Helliwell et al., 2021). These advances will now allow detailed examination of diatom signalling in response to range of stimuli, including temperature.

In this study we set out to examine the ability of diatoms to sense short-term changes in temperature. In particular, we examined whether the well-characterised [Ca^2+^]_cyt_ elevations observed in animal and plant cells in response to rapid changes in temperature were conserved in diatoms. Using the model species *P. tricornutum* and *T. pseudonana*, which can both inhabit coastal environments that experience variable temperature regimes (De Martino et al., 2007; Alverson et al., 2011), we found that diatoms consistently exhibit a [Ca^2+^]_cyt_ elevation in response to cold shock, but do not exhibit [Ca^2+^]_cyt_ elevations in response to elevated temperature. We did not find a requirement for cold shock-induced Ca^2+^ signalling in increasing tolerance to low temperatures, but found that cold shock increases resistance to simultaneous hypo-osmotic shocks, suggesting that integration of multiple signalling inputs may contribute to an enhanced ability to respond to these environmental stimuli.

## Methods

### Recording of rockpool temperature

Temperature data was recorded using a 27 mm Envlogger v2.4 (ElectricBlue, Porto, Portugal) encased in acrylic resin, recording in 30 minute intervals with a resolution of 0.1 °C. The Envlogger was secured to the substrate using Z-Spar A-788 epoxy resin roughly 3 cm below the surface waters of a shallow midshore rockpool measuring approximately 8 cm deep at Looe Hannfore, Cornwall, UK (50.3411, -4.4598) from 1/7/2019 to 7/7/2019.

### Strains and culturing conditions

The wild type *P. tricornutum* strain used in this study was CCAP 1055/1 (Culture Collection of Algae and Protozoa, SAMS, Scottish Marine Institute, Oban, UK). A *P. tricornutum* strain transformed with the R-GECO1 Ca^2+^ biosensor (PtR1) and the three *eukcata1* knock-out strains in this line (labelled A3, B3 and B6) were generated as described previously(Helliwell et al., 2019). The *T. pseudonana* strain expressing the R-GECO1 biosensor (TpR1) was generated as described in Helliwell et al (2021). Cultures were maintained in natural seawater with f/2 nutrients(J C Lewin and Guillard, 1963; Guillard, 1975); modified by the addition of 106 μM Na_2_SiO_3_.5H_2_O and the exclusion of vitamins (*P. tricornutum* only). For imaging experiments, cells were acclimated to an artificial seawater (ASW) medium for minimum 10 days prior to analysis. ASW contained 450 mM NaCl, 30 mM MgCl_2_, 16 mM MgSO_4_, 8 mM KCl, 10 mM CaCl_2_, 2 mM NaHCO_3_, 97 µM H_3_BO_3_, f/2 supplements and 20 mM HEPES (pH 8.0). Cultures were grown at 18 °C with a 16:8 light/dark cycle under illumination of 50 µmol m^-2^ s^-1^.

### Epifluorescence imaging of R-GECO1 fluorescence

500 µL of cell culture was added to a 35 mm microscope dish with glass coverslip base (In Vitro Scientific, Sunnyvale, CA, USA) coated with 0.01% poly-L-lysine (Merck Life Science UK, Gillingham, Dorset) to promote cell adhesion to the glass surface. Cells were allowed to settle for 5-20 minutes at room temperature (RT) under light. R-GECO1 was imaged using a Leica DMi8 inverted microscope (Leica Microsystems, Milton Keynes, UK) with a 63x 1.4NA oil immersion objective, using a Lumencor SpectraX light source with a 541-551 nm excitation filter and 565-605 nm emission filter. Images were captured with a Photometrics Prime 95B sCMOS camera (Teledyne Photometrics, Birmingham, UK). Images were captured at 3.33 frames per second using Leica application suite X-software v.3.3.0.

### Administration of temperature shocks to cells in the imaging setup

The dish was perfused with ASW without f/2 nutrients at a standard flow rate of 16 mL min^-1^. To achieve rapid changes in temperature in the dish, the perfusion was switched between solutions of different temperature to achieve target temperatures of approximately 10, 22 or 30 °C respectively. Actual dish temperature was recorded using a Firesting micro optical temperature sensor (Pyroscience GmbH, Aachen, Germany). For the majority of experiments, cells were perfused with warmer media (dish temperature 30 °C) for 1 minute prior to application of the cooling shock (these conditions reflect those observed in the rockpool observations). The perfusion flow rate was altered to achieve different temperature change rates. As cooling rate was not linear, the maximum cooling rate was defined as the largest decrease temperature within a one second period.

### Application of inhibitors and elicitors

External Ca^2+^ was removed by perfusion with ASW without CaCl_2_ containing 200 μM EGTA. Ruthenium red (RR) was added to cells at a final concentration of 10 µM 5 minutes prior to cold shock treatment. Menthol was prepared as a 1 M stock solution in DMSO and used at concentration of 1 mM, resulting in a final DMSO concentration of 0.1%.

### Processing of imaging data

Images were processed using LasX software (Leica). The mean fluorescence intensity within a region of interest (ROI) over time was measured for each cell by drawing an ROI encompassing the whole cell. Background fluorescence was subtracted from all cellular F values. The change in the fluorescence intensity of R-GECO1 was then calculated by normalizing each frame by the initial value (F/F_0_). [Ca^2+^]_cyt_ elevations were defined as any increase in F/F_0_ above a threshold value (1.5). The duration of a [Ca^2+^]_cyt_ elevation was defined as the peak width at half maximal amplitude. To visualise the spatial distribution of a [Ca^2+^]_cyt_ elevation, each frame was divided by a corresponding background image generated by applying a rolling median (30 frames) to the image-series (Image J). The resultant time series images were pseudo-coloured to indicate changes in fluorescence.

### Statistical analysis

Graphs and statistical analyses were performed using Sigmaplot v14.0 (Systat Software, Slough, UK). Error bars represent standard error of the mean. Unless indicated otherwise, imaging experiments were repeated three times with independent cultures on different days to ensure reproducibility of the response.

Normal distribution of respective datasets was tested using Shapiro-Wilk’s normality test. When passed, statistical analysis of datasets with two groups were done with Student’s t-test, and when not passed with Mann-Whitney’s rank sum test. Statistical analysis of datasets with more than two groups were performed using an ANOVA followed by a Holm-Sidak post-hoc test when the normality test was positive. When the normality test was negative, Kruskal-Wallis’ 1-way Analysis of Variance on Ranks was used instead. All statistical tests were performed with Sigmaplot v14.0.3.192 (Systat software Inc).

### Growth at different temperatures after a cold shock

For the growth curves, cells were grown to mid exponential phase (2.73 × 10^6^ cells mL^-1^). The culture was divided into 10 mL aliquots and cells were pelleted by centrifugation (4000 x *g* at 18 °C). Cells were washed in 40 mL ASW +/- Ca^2+^ and pelleted again by centrifugation. Cells were then resuspended in their respective treatments (20 mL of ASW +/- Ca^2+^ at 18 °C or 4 °C to administer a rapid cold shock). After 10 minutes, 2 mL of each culture was used to inoculate culture vessels containing 18 mL of standard F/2 media (approx. starting density of 6.8 × 10^4^ cells mL^-1^) and cultures were grown at 18 °C or 4 °C.

### Cell survival during hypo-osmotic shock

To examine the effect of temperature on cell viability during hypo-osmotic shock, 10 mL of a late log phase culture (6 × 10^6^ cells mL^-1^) were pelleted by centrifugation (4000 x *g* at 18 °C). Cells were washed twice with 10 mL ASW +/-Ca^2+^ and 250 μL aliquots were taken. To apply the hypo-osmotic and cold shock treatments, 750 µL of ASW (+/- Ca^2+^) or deionised water at two different temperatures (20 °C or 4 °C) were added to each tube. Addition of water results in a severe hypo-osmotic osmotic shock (final concentration 25% ASW) simultaneously with the temperature shock. The cells were then incubated at their respective temperatures for 3 minutes prior to addition of 5 μM Sytox Green (Thermo Fisher Scientific, Loughborough, UK). All treatments were then incubated at 20 °C 15 min in darkness. Cell viability was measured with a LUNA-FL(tm) Dual Fluorescence Cell Counter (Logos Biosystems, Villeneuve d’Ascq, France) to count live (displaying red chlorophyll fluorescence) versus dead cells (Sytox Green stain) with following settings: Excitation intensity green = 11, red = 7, count threshold for both = 3.

### Quantification of K+ efflux in P. tricornutum populations using K+ -selective microelectrodes

K^+^ microelectrodes were fabricated as described previously (Helliwell et al., 2021). Clark GC-1.5 borosilicate glass capillaries (Harvard Apparatus, Cambridge, UK) were pulled to a fine point using a P-97 pipette puller (Sutter, Novato, CA). The pipette tips were then gently broken to produce a diameter of ca 10-20 µm. The capillaries were silanised by exposure to N,N-dimethyltrimethylsilylamine (TMSDMA) vapour at 200 °C for 20 minutes within a closed glass Petri dish. The K^+^ microelectrodes were prepared by introducing K^+^ ionophore I (Sigma Aldrich, Gillingham, Dorset, UK) into the pipette tip by suction. Pipettes were then back-filled with the filling solution (100 mM NaCl, 20 mM HEPES pH 7.2 and 10 mM NaOH). The reference electrode was filled with 3 M KCl and data were recorded using an AxoClamp 2B amplifier, with pClamp v10.6 software (Molecular Devices, CA, USA). Each K^+^ microelectrode was calibrated using a two-point calibration with standard KCl solutions. The mean slope of the calibrated electrodes was 53.0 ± 1.3 mV per decade (±SE).

For the measurements, 10 mL of *P. tricornutum* cells from exponential culture containing 10^6^- 10^7^ cells mL^-1^ were centrifuged at 4000 rpm for 10 min and re-suspended in 1 mL of ASW. The cells were then allowed to settle on a poly-L-lysine coated microscope dish. Cells were perfused with ASW or ASW -Ca^2+^ (0 μM Ca^2+^ + 100 µM EGTA) and cold shocks were applied as described for the microscopy observations. Control experiments were performed in the absence of *P. tricornutum* cells to ensure that the change in temperature did not alter the performance of the K^+^ microelectrodes.

## Results

### Rapid changes in temperature in inter-tidal environments

*P. tricornutum* was first isolated from a tidal pool in the UK and has since been identified in a range of coastal and brackish habitats (De Martino et al., 2007). To assess the dynamic temperature regimes potentially experienced by intertidal diatoms, we measured the temperature of a tidal pool located on the upper region of a rocky shore (South Cornwall, UK) over a 7 d period during July (UK summer). Temperatures within the pool were very stable around 15°C during immersion at high tide (Fig 1). However, at low tides temperatures in the exposed tidal pool rose significantly during the day (up to 30 °C) and decreased at night (to 12 °C), before being rapidly restored to the bulk seawater temperature by the immersion of the pool at high tide. These data illustrate that diatoms inhabiting intertidal environments in temperate regions will regularly experience periods of significant warming followed by rapid cooling. The fluctuations in temperature are likely to be even greater in smaller volumes of water, such as the surface of estuarine mudflats or very shallow pools.

**Figure 1:**
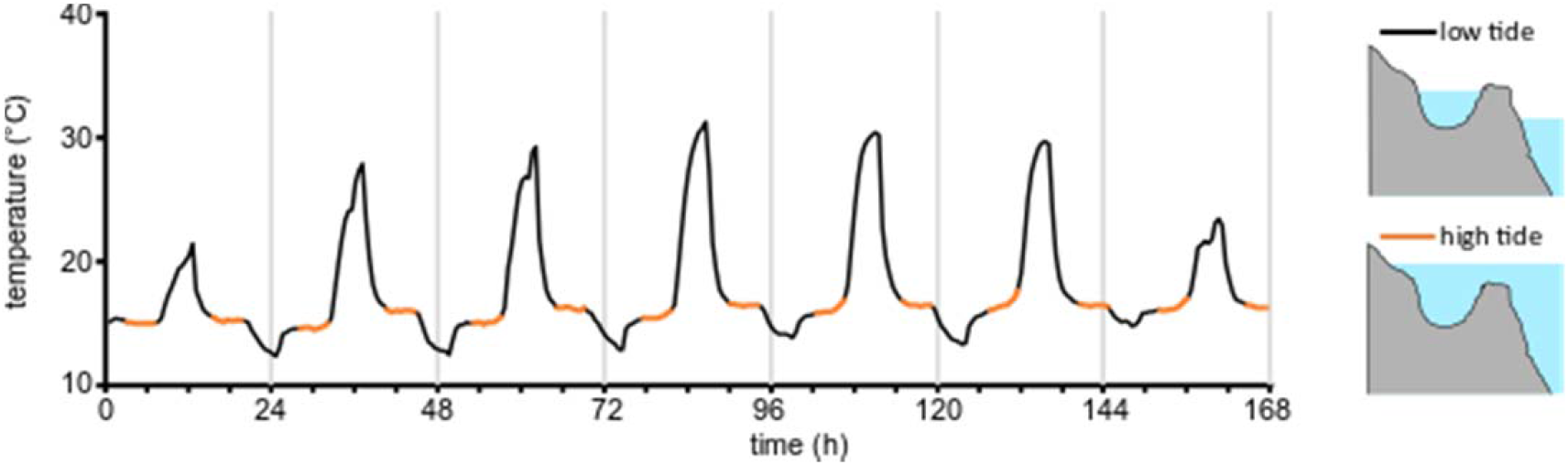
Temperature fluctuations in the inter-tidal zone. An example of temperature fluctuations measured in a temperate coastal rock pool (Looe, Cornwall, UK) over the course of seven days in summer (01/07/2019-07/07/2019). Orange colour indicates periods at which the pool was immersed by the high tide (approx. duration of immersion 5 h). Significant excursions from the sea temperature occur when the rock pool is isolated from the bulk seawater at low tide (black traces). Rapid cooling (30 °C to 15 °C) occurs when the incoming tide reaches the pool.

### Calcium signalling in response to changes in temperature

*P. tricornutum* cells expressing the R-GECO1 Ca^2+^ biosensor were perfused with seawater at high or low target temperatures (30 °C or 12 °C). Note that actual temperatures in the perfusion dish differed by ± 2 °C from these target temperatures due to equilibration of the small volume of warm or cold perfusate with room temperature. Actual dish temperatures were therefore recorded and are displayed for all experiments. We routinely observed a single transient [Ca^2+^]_cyt_ elevation in cells exposed to a cold shock from 30 °C to 12 °C (97 % cells, n=63) (Fig 2A). In contrast, cells exposed to a rapid rise in temperature from 12 °C to 30 °C did not show [Ca^2+^]_cyt_ elevations (Fig. 2A). No [Ca^2+^]_cyt_ elevations were observed in cells perfused with these solutions after they had been equilibrated to room temperature, indicating that the act of switching between the perfusion solutions does not contribute to the signalling responses (Fig. 2A). Analysis of the spatial characteristics of cold-shock induced [Ca^2+^]_cyt_ elevations indicated that many initiate at the apex of the cell and propagate to the towards the central region (Fig 2B), in a manner similar to those induced by mild hypo-osmotic shock (Helliwell et al., 2021). This suggests that the apices of the cell may play an important role in sensing the temperature changes. Cells exposed to a second cold shock two minutes after a previous cold shock demonstrated [Ca^2+^]_cyt_ elevations with no significant attenuation in amplitude, although the percentage of cells responding was slightly lower (97% to 81 % of cells, n=63) (Supplementary Fig S1).

**Figure 2:**
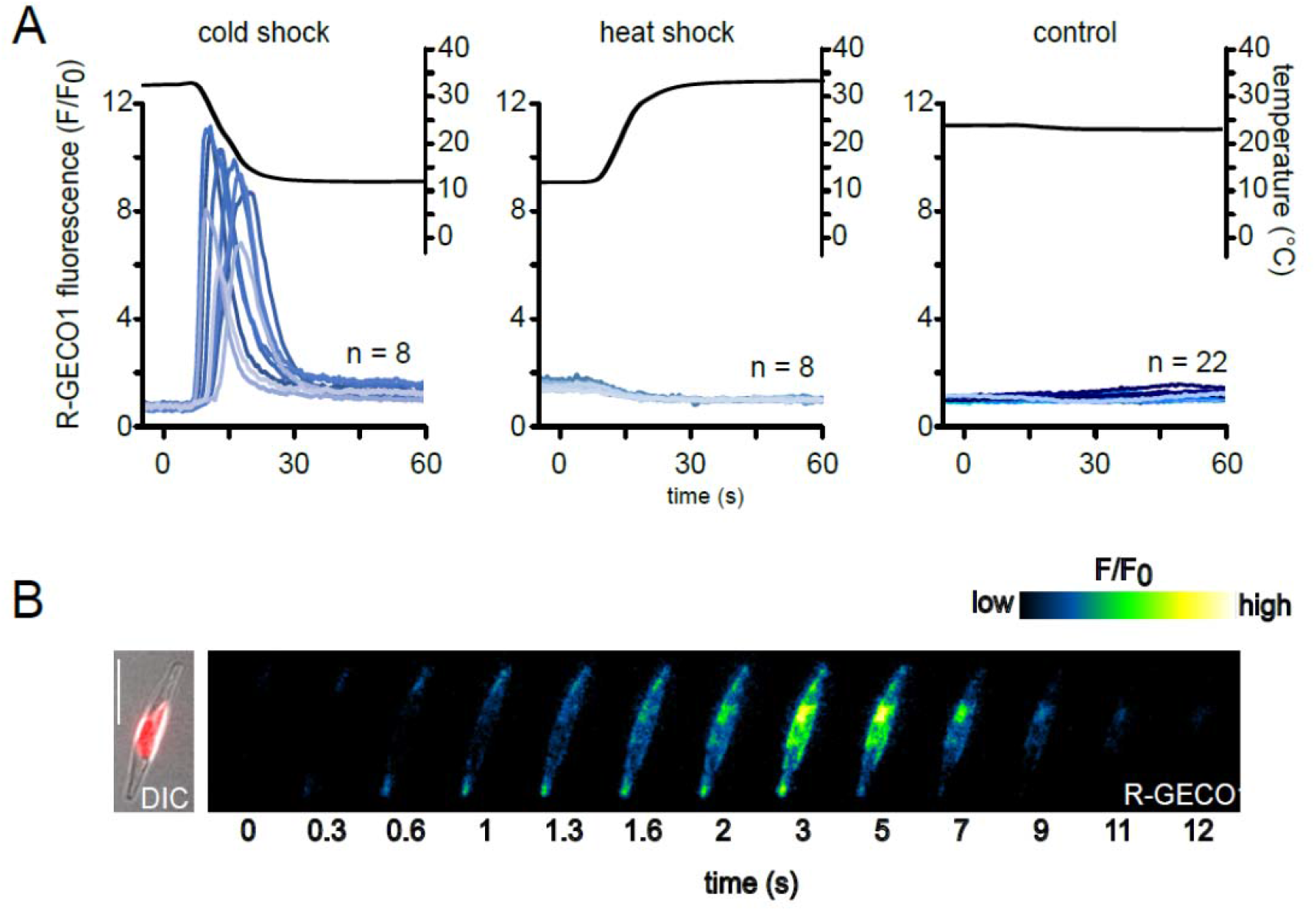
*P. tricornutum* exhibits cytosolic Ca^2+^([Ca^2+^]_cyt_) elevations in response to rapid cooling. **A)** Eight representative fluorescence ratio traces (F/F_0_, blue lines) of *P. tricornutum* cells expressing R-GECO1 representing changes in cytosolic Ca^2+^. Cells were perfused with ASW of different temperatures to cause rapid temperature shifts (black line). Cold shock 30 °C to 12 °C, heat shock 12 °C to 30 °C or control 22 °C to 22 °C. **B)** False colour images of a PtR1 cell exhibiting a [Ca^2+^]_cyt_ elevation in response to cold shock. The temperature decrease begins at t = 0 s. Note that the [Ca^2+^]_cyt_ elevations initiate at the tips of the cell and spread towards the central region. Left panel indicates a DIC image overlaid with chlorophyll autofluorescence. Bar represents 10 µm.

The [Ca^2+^]_cyt_ elevations observed during cold shock were represented by a >10-fold increase in R-GECO1 fluorescence. Assuming a K_d_ of 480 nM for R-GECO1 and comparison with published maximum F/F_0_ values (Zhao et al., 2011), we estimate that [Ca^2+^]_cyt_ elevations reach concentrations in the μM range, which are likely to be physiologically significant. In addition to these large increases in fluorescence that are attributed to [Ca^2+^]_cyt_ elevations, much smaller changes in the baseline fluorescence of each cell could be observed following changes in temperature (increasing with low temperature and decreasing with high temperature, Supplemental Fig S1). These minor changes most likely represent temperature-dependent changes in R-GECO1 fluorescence emission (Ohkura et al., 2012) rather than actual changes in resting Ca^2+^ concentration. Therefore, only the substantial transient increases in fluorescence (F/F_0_>1.5) representing large [Ca^2+^]_cyt_ elevations were analysed further.

[Ca^2+^]_cyt_ elevations were also observed when a cold shock (30°C to 12 °C) was applied to cells held at 22 °C, rather than 30 °C, indicating that the cold shock response was not a consequence of prior warming of the cells (Supplemental Fig S2). The percentage of cells responding to cold shock was lower in cells held at 22 °C compared to cells held at 30 °C, although this may also be influenced by the lower maximum rate of cooling at 22 °C.

### Rapid cooling is required to elicit a [Ca^2+^]_cyt_ elevation

We therefore examined the nature of the temperature change required to elicit [Ca^2+^]_cyt_ elevations, by manipulating the flow rate of the perfusion to vary the rate of cooling. Rapid cooling (2.5 °C s^-1^) resulted in [Ca^2+^]_cyt_ elevations in 100% of cells examined (n=45), whereas only 7% of cells exhibited a [Ca^2+^]_cyt_ elevation at a cooling rate of 0.4 °C s^-1^ (n=45) (Fig. 3A-B). The amplitude of the [Ca^2+^]_cyt_ elevations in responding cells closely correlated with the cooling rate, with much larger [Ca^2+^]_cyt_ elevations observed at rapid cooling rates (Fig. 3C). Examination of a broader range of cooling rates indicated that a cooling rate greater than 1 °C s^-1^ was required to elicit [Ca^2+^]_cyt_ elevations in 50% of the population (Fig. 3D). These data suggest that the cold shock-induced [Ca^2+^]_cyt_ elevations can therefore relay information relating to the nature of the stimulus both in terms of the number of cells responding and the nature of the [Ca^2+^]_cyt_ elevation itself.

**Figure 3:**
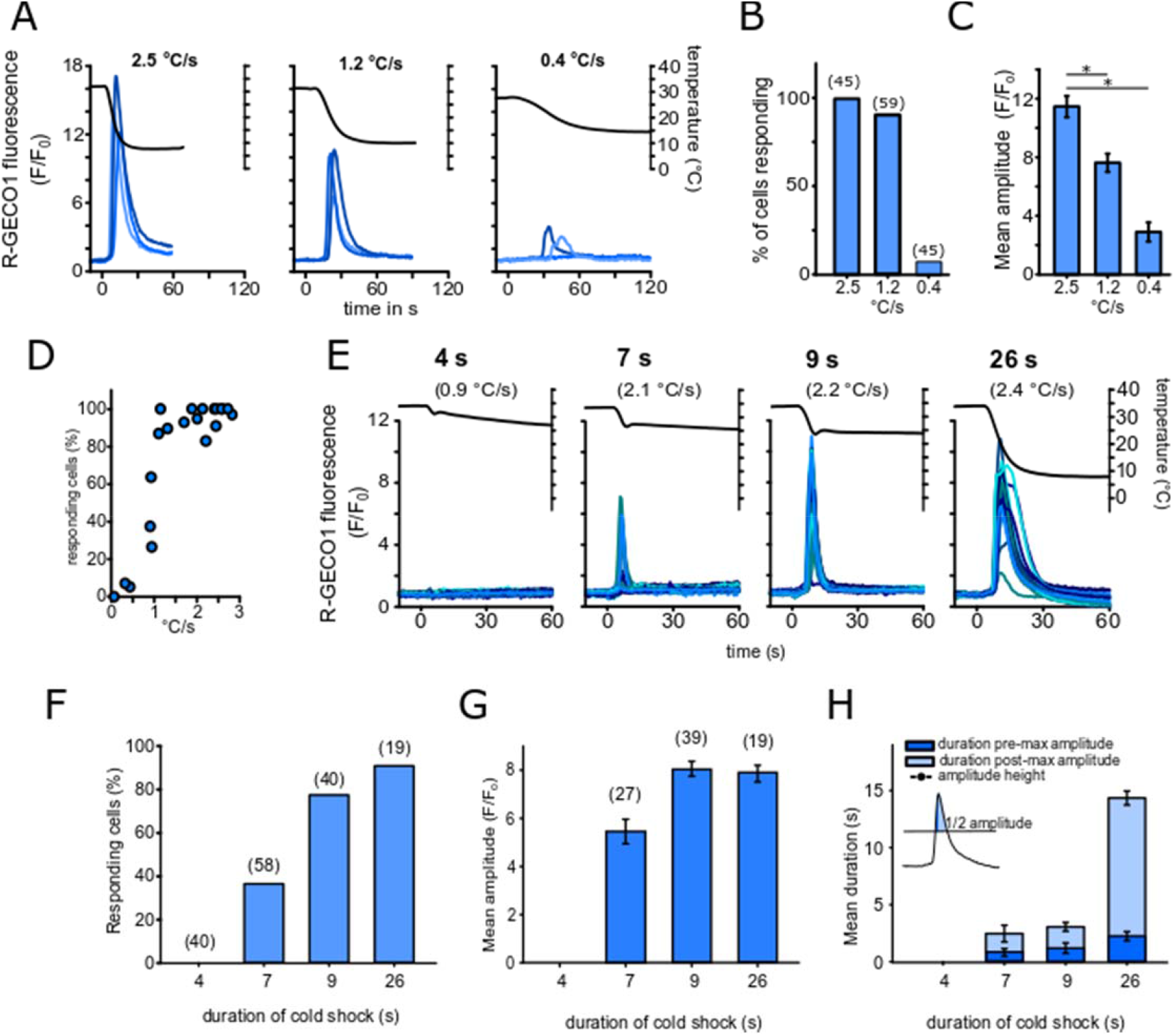
The cold shock Ca^2+^response depends on the rate of change of temperature. **A)** R-GECO1 fluorescence in *P. tricornutum* in response to cold shock administered at different cooling rates. As cooling rates were non-linear the maximal cooling rate for each treatment was calculated for comparisons. Three representative traces are shown. **B)** The percentage of cells exhibiting a [Ca^2+^]_cyt_ elevation (F/F_0_>1.5) at different cooling rates. Total number of cells examined are shown in parentheses, from a minimum of two separate experimental treatments. **C)** Mean maximal amplitude of [Ca^2+^]_cyt_ elevations from responsive cells in (B). * indicates a significant difference (One way ANOVA on Ranks P>0.001, Dunn’s post-hoc test P>0.001). n=31,30 and 3 for 2.5, 1.2 and 0.4 °C s^-1^ respectively. Error bars = SE. **D)** Percentage of cells responding to cold shock with a [Ca^2+^]_cyt_ elevation across a broader range of maximum cooling rates. The data represent 21 independent experiments, with a mean of 38 cells examined for each data point (minimum 12, maximum 123 cells). **E)** [Ca^2+^]_cyt_ elevations in response to different durations of cooling applied with a constant flow rate (16 mL min^-1^). 20 representative traces from PtR1 cells are shown, with greater [Ca^2+^]_cyt_ elevations observed under increasing durations of cold shock. The maximum rate of temperature decrease (ΔT s^-1^) is shown in parantheses. Data for 4, 7, 9 and 26 s of cold shock duration were compiled from 2, 3, 2 and 1 individual experiments, respectively. **F)** The percentage of cells exhibiting a [Ca^2+^]_cyt_ elevation in response to cold shock for the experiment described in (E).**G)** Mean maximal amplitude of [Ca^2+^]_cyt_ elevations in response to cold shock for the responding cells shown in (F). **H**) Duration of [Ca^2+^]_cyt_ elevations (shown as full width at half maximum amplitude) in relation of the duration of cold stimulus. The duration of [Ca^2+^]_cyt_ elevations is greatest at the 26 s cold shock. The duration is divided into a pre- and post-maximal amplitude component to show that the post-maximal amplitude (tail) components of the [Ca^2+^]_cyt_ elevation is greatly extended under the 26 s cold shock.

As very low perfusion rates also resulted in a lower overall decrease in temperature (due to equilibration of the perfusate with room temperature), we next examined the absolute temperature decrease required to initiate signalling. Cells were perfused at 30 °C for 1 minute and then perfused at a constant flow rate with cold ASW (4 °C) for different durations to vary the decrease in temperature whilst maintaining similar rates of cooling. A very brief perfusion (4 s) lowered the temperature by 2.4 ±0.6 °C but did not induce [Ca^2+^]_cyt_ elevations (Fig 3E-F). However, the rate of cooling in this treatment was considerably lower than the other treatments, due to buffering of the temperature by the residual volume within the perfusion dish (1 mL). Perfusions of a longer duration (7-26 s) resulted in a consistent cooling rate of 2.1-2.4 °C s^-1^. A temperature decrease of 8.8 ± 0.4 °C induced a [Ca^2+^]_cyt_ elevation in 38.4% of cells (n=126), whereas greater decreases in temperature resulted in [Ca^2+^]_cyt_ elevations in nearly all cells (Fig. 3E-F). The amplitude and duration of [Ca^2+^]_cyt_ elevations increased with the greater duration of the temperature decrease (Fig 3G-H). A cooling duration of 26 s did not increase the amplitude of the [Ca^2+^]_cyt_ elevation beyond those observed at 9 s, but greatly increased the duration of the [Ca^2+^]_cyt_ elevation (Fig 3G-H). Taken together, our results show that the cooling rate and the duration of the cold shock influence the amplitude and duration of the [Ca^2+^]_cyt_ elevation and the percentage of cells responding.

### The cold shock response is conserved in the centric diatom *T. pseudonana* but displays different characteristics

*T. pseudonana* is a planktonic centric diatom found in marine, estuarine and freshwater environments (Alverson et al., 2011), where it is also likely to be exposed to significant changes in temperature. We found that *T. pseudonana* cells expressing the R-GECO1 biosensor exhibited [Ca^2+^]_cyt_ elevations in response to cold shock, with the amplitude of these elevations also dependent on the rate of temperature decrease (Fig. 4A-B). As in *P. tricornutum*, a control perfusion using ASW media equilibrated to room temperature did not induce [Ca^2+^]_cyt_ elevations (Fig. 4C). The cold induced [Ca^2+^]_cyt_ elevations in *T. pseudonana* were of a lower amplitude but longer duration than those observed in *P. tricornutum*. The percentage of *T. pseudonana* cells responding to cold shock was also considerably lower than *P. tricornutum* (19 % vs. 81 %, respectively), although variable levels of expression of R-GECO1 in *T. pseudonana* likely prevented detection of [Ca^2+^]_cyt_ elevations in all cells within a field of view (Helliwell et al., 2021) (Fig. 4D-F). Taken together, these findings suggest that cold shock-induced [Ca^2+^]_cyt_ elevations are exhibited by both pennate and centric diatom lineages and may therefore represent a conserved mechanism in many diatom species.

**Figure 4:**
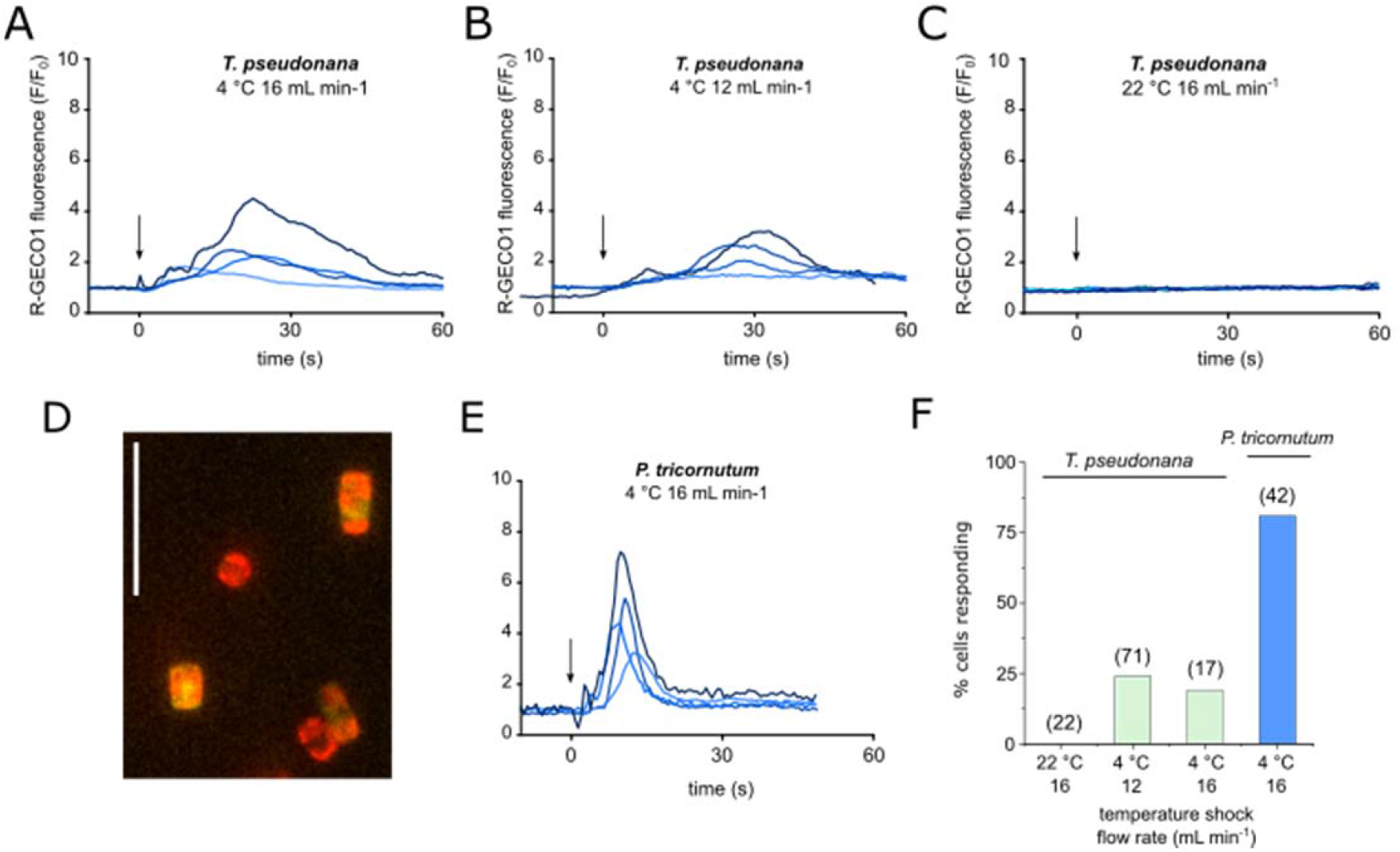
*Thalassiosira pseudonana* also shows cold-induced [Ca^2+^]_cyt_ elevations. **A)** Fluorescence ratio of *T. pseudonana* cells expressing cytosolic R-GECO1 in response to a cold shock (from 30 to 10 °C). For these experiments the temperature in the dish was not monitored, so perfusion flow rate is shown to indicate rate of cold shock. Arrow indicates onset of cold stimulus. Four representative traces are shown. **B)** As in (A) but at a slower flow rate. **C)** Treatment control using perfusion of ASW at room temperature. **D)** Fluorescence image of *T. pseudonana* cells expressing R-GECO1 (yellow) overlaid with chlorophyll autofluorescence (red). Scale bar represents 20 µm. **E)** *P. tricornutum* cold shock response under identical treatment as in **A** for comparison. **F)** Percentage of cells exhibiting [Ca^2+^]_cyt_ elevations. Values in parentheses denote n.

### Cellular mechanisms underlying the cold shock response

We next examined the cellular mechanisms responsible for cold shock Ca^2+^ signalling in *P. tricornutum*. Removal of external Ca^2+^ by perfusion of PtR1 cells with cold Ca^2+^-free ASW completely abolished the [Ca^2+^]_cyt_ elevations (Fig. 5A). Restoration of external Ca^2+^ to cooled cells did not induce a [Ca^2+^]_cyt_ elevation. However, when these cells were subsequently warmed to 30 °C and then cooled, [Ca^2+^]_cyt_ elevations were observed in the majority of cells. Thus, the generation of cold-induced [Ca^2+^]_cyt_ elevation depends on the presence of external Ca^2+^, and the [Ca^2+^]_cyt_ elevation is triggered by the rapid drop in temperature rather than low absolute temperature itself.

**Figure 5:**
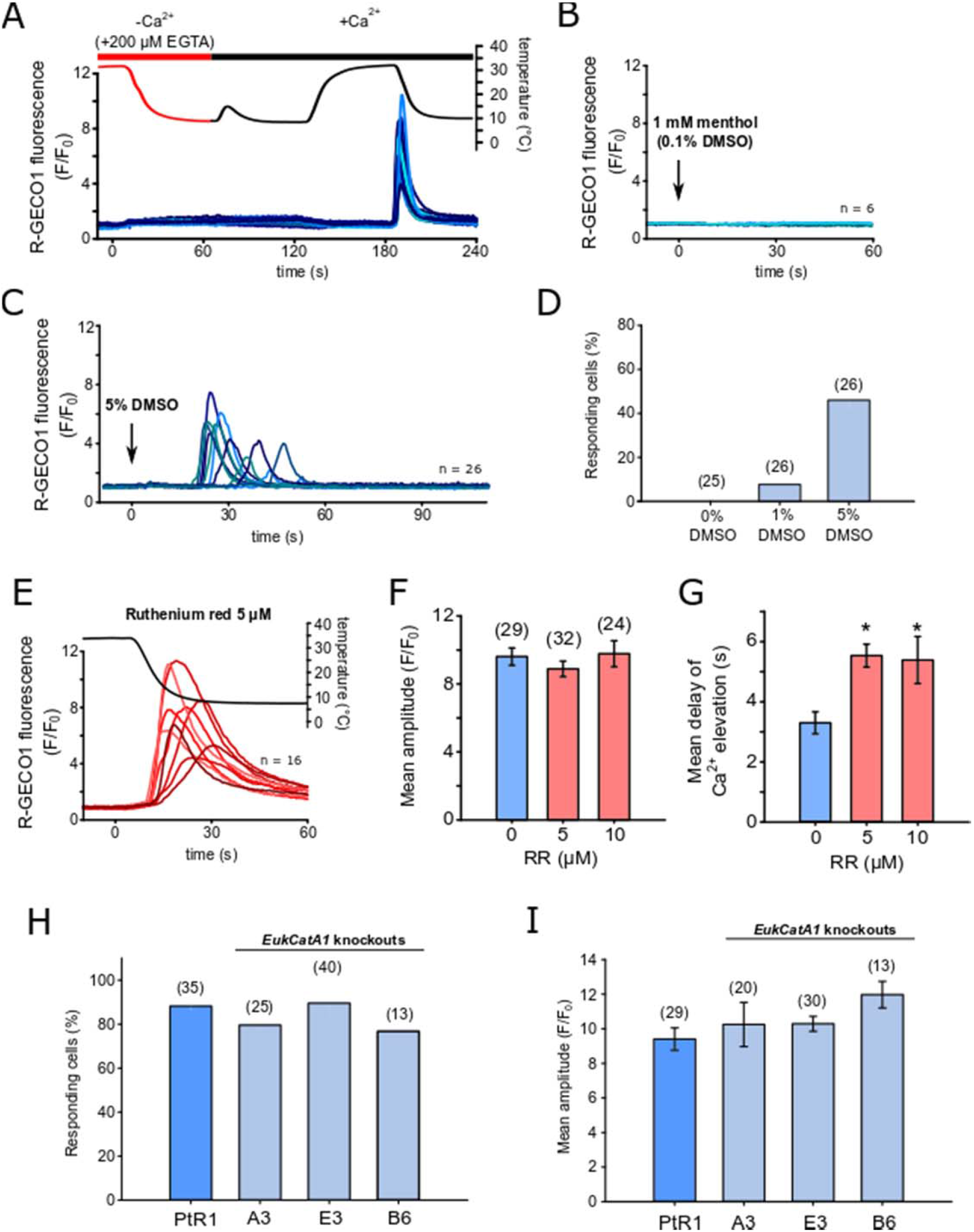
Cellular mechanisms of cold shock induced [Ca^2+^]_cyt_elevations. **A**) R-GECO1 fluorescence ratio (F/F_0_) from a cold shock applied to PtR1 cells using ASW without Ca^2+^ + 200 μM EGTA (Methods). No [Ca^2+^]_cyt_ elevations can be observed during the cold shock or when Ca^2+^ was restored to cold cells (perfused with cold ASW with Ca^2+^). However, [Ca^2+^]_cyt_ elevations were observed during a subsequent cold shock applied with standard ASW (i.e. with 10 mM CaCl_2_). Note the minor temperature increase at 70 s is due to a slight warming of cold ASW+Ca^2+^ media in the perfusion system. 23 representative traces are shown, three additional experiments were performed with identical results. **B)** PtR1 fluorescence in response to ASW containing 1 mM menthol (including 0.1% DMSO as solvent carrier). Six representative traces are shown. **C)** R-GECO1 fluorescence ratio of PtR1 cells perfused with ASW + 5% DMSO. **D)** Percentage of cells exhibiting [Ca^2+^]_cyt_ elevations in response to DMSO. **E)** The effect of cold shock on PtR1 cells pre-treated with the Ca^2+^ channel blocker ruthenium red (5 μM final, 5 min pre-incubation). 16 representative traces are shown. **F)** Mean amplitude (±SE) of responding cells treated with ruthenium red (5 μM or 10 μM) compared to untreated control cells. No significant differences were observed (one-way ANOVA). **G)** Mean timing (±SE) of the maximal amplitude of [Ca^2+^]_cyt_ elevations for cells treated with ruthenium red (P = <0.01, one-way ANOVA, Holm-Sidak post-hoc test). **H)** Percentage of cells showing [Ca^2+^]_cyt_ elevations in response to cold shock in control and three independent *eukcata1* mutant strains (30 to 10 °C). Data represent a minimum of two independent experiments per strain (one-way ANOVA). **I)** Mean maximal amplitude (±SE) of [Ca^2+^]_cyt_ elevations in *eukcatA1* mutants in response to cold shock. No significant differences were observed (one-way ANOVA on Ranks).

*P. tricornutum* lacks cyclic-gated nucleotide channels, which are important for thermal sensing in plants, although it does possess multiple TRP channels (Verret et al., 2010). The temperature-sensitive TRPM8 channel in animal cells is responsible for cold induced [Ca^2+^]_cyt_ elevations and can be activated directly by the plant secondary metabolite, menthol (Yin et al., 2018). Perfusion of PtR1 cells with 1 mM menthol did not elicit [Ca^2+^]_cyt_ elevations, indicating that this ligand is likely specific to the ion channels involved in animal cold signalling (Fig. 5B). In plant and fungal cells, cold shock induced [Ca^2+^]_cyt_ elevations have been studied through the application of dimethyl sulfoxide (DMSO), which is proposed to mimic cold-induced membrane rigidification (Furuya et al., 2014). DMSO elicited [Ca^2+^]_cyt_ elevations in a dose-dependent manner in *P. tricornutum*, with 8% and 50% of cells exhibiting Ca^2+^ elevation in response to addition of 1 % and 5% DMSO respectively (n = 24, 25) (Fig.5C-D). Ruthenium red (RR) is a non-selective Ca^2+^ channel blocker shown to affect numerous TRP channels including the cold sensitive TRPA1 channel (Andrade et al., 2008; Silva et al., 2015; Christensen et al., 2016). RR also inhibits [Ca^2+^]_cyt_ elevations in *P. tricornutum* induced by the resupply of phosphate to phosphate–limited cells, but does not inhibit [Ca^2+^]_cyt_ elevations caused by hypo-osmotic shock (Helliwell et al., 2021; Helliwell et al., 2021). Pre-treatment of PtR1 cells for 5 mins with 5-10 μM RR did not significantly reduce the amplitude of cold-induced Ca^2+^ elevations (Fig 5E-F). However, RR treated cells exhibited a significantly slower response time than non-treated control cells (defined as time from stimulus to the initial elevation above the threshold of F/F_0_>1.5) (Fig. 5G), indicating that whilst RR does not prevent the cold shock response, it may partially inhibit a component of the signalling pathway.

*P. tricornutum* contains three EukCatA channels, which represent a novel class of voltage-gated Na^+^/Ca^2+^ channel related to single-domain voltage-gated Na^+^ channels in bacteria (BacNav) (Helliwell et al., 2019). As BacNav channels are temperature sensitive (Arrigoni et al., 2016), we examined whether *P. tricornutum* e*ukcatA1* knockout mutants exhibited an altered response to cold shock. The percentage of responding cells and the mean maximal amplitude of the [Ca^2+^]_cyt_ elevations did not differ significantly from control PtR1 cells (Fig. 5H-I), indicating that EukCatA1 is not required for cold shock induced Ca^2+^ signalling. Further experiments will be needed to determine whether other candidate ion channels, such as the other EukCatA channels or the diatom TRP channels, contribute to this signalling response.

### The cold shock response is not required for growth at low temperatures

We next examined whether the cold shock Ca^2+^ signal was required for *P. tricornutum* cells to survive following a cold shock. We applied cold shocks in the presence and absence of external Ca^2+^ and then monitored their physiology and subsequent growth at 4 °C or 18 °C in the presence of Ca^2+^ (Fig 6A). While cells grew significantly more slowly at 4 °C versus 18 °C, there was no significant impact of inhibiting Ca^2+^ signalling during cold shock on the ability of cold-shocked cells to grow at either 4 °C or 18 °C (Fig 6B).

**Figure 6:**
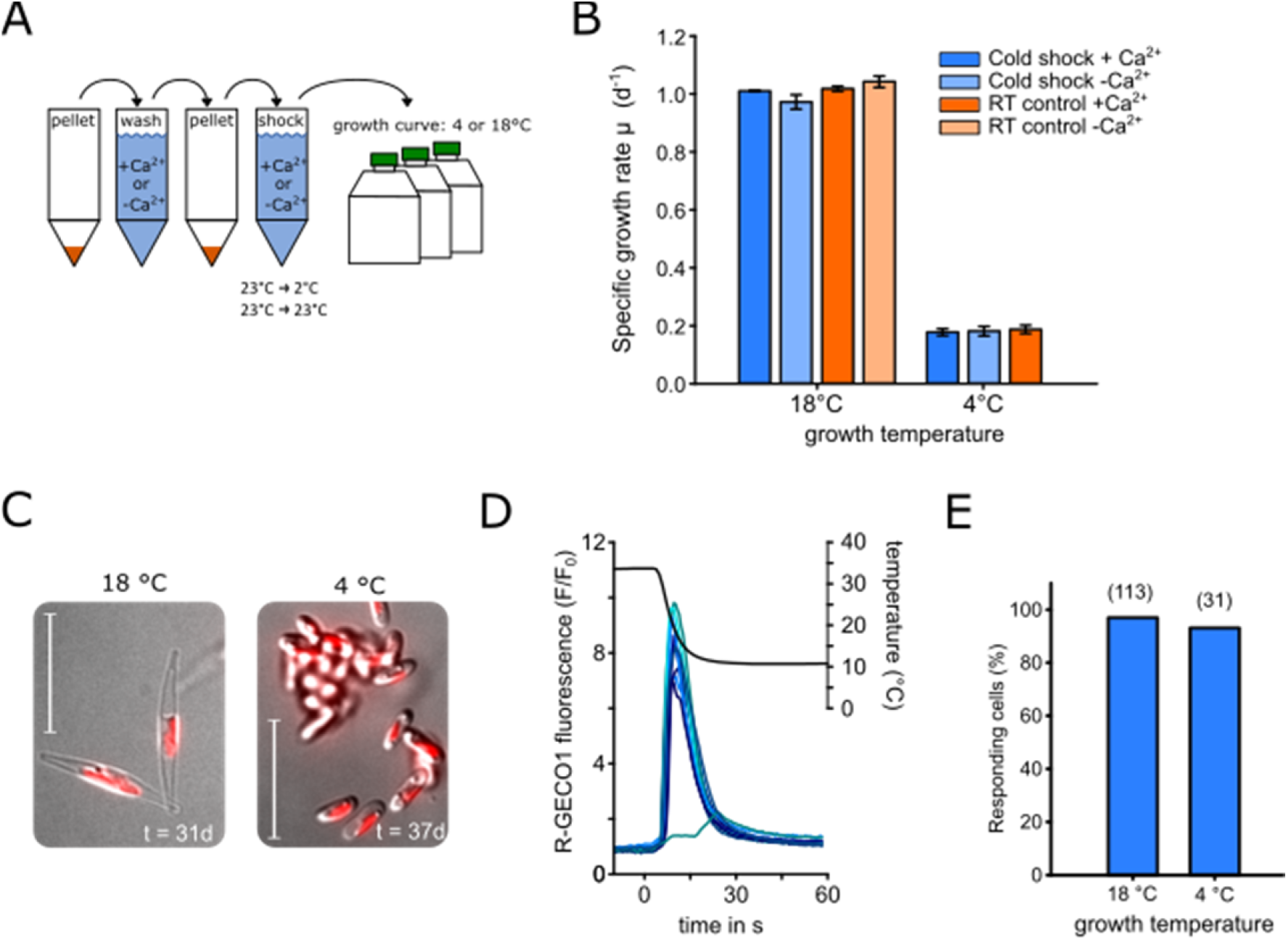
The role of the cold shock response in cold tolerance. **A)** Schematic diagram showing the workflow for an experiment examining the impact of Ca^2+^-dependent cold shock signalling on *P. tricornutum* cold tolerance. Cells were harvested and washed in ASW containing 10 mM Ca^2+^ or no Ca^2+^ (ASW –Ca^2+^ +200 µM EGTA). Cells were pelleted once again and exposed to cold shock with or without Ca^2+^. Cells were then grown at control (18 °C) and cold (4 °C) conditions in standard ASW (i.e. with 10 mM CaCl_2_) to examine cold tolerance. **B)** Growth rate of *P. tricornutum* cultures after cold shock treatment. Mean specific growth rates were calculated from day 0-5 and 12-30 for 18 °C and 4 °C, respectively. Note that for growth at 4 °C, a no shock (RT) control in the absence of Ca^2+^ was not included. No significant differences were observed between treatments at each temperature (one-way ANOVA). Error bars represent SE. **C)** DIC images of PtR1 cells grown at 18 or 4 °C. Oval cells predominate in cells grown at 4 C for extended periods. Red = chlorophyll auto-fluorescence, bar = 20 µm. **D)** Cold-acclimated PtR1 cells still exhibit a response to cold shock. Representative R-GECO1 fluorescence ratio traces from PtR1 cells grown at 4 °C for 4 days. Cells were briefly warmed to 30 °C before a cold shock was applied. **E)** The percentage of cells responding to cold shock as function of acclimation temperature. The results were generated from four separate experiments with maximum temperature drop-rates between 2.2 - 3.2 C° s^-1^.

Growth of *P. tricornutum* at 4 °C promoted the accumulation of the oval morphotype, as reported previously (De Martino et al., 2011) (Fig 6C). Cells acclimated to low temperatures may therefore undergo physiological changes that render them less sensitive to rapid cooling. However, fusiform cells grown at 4 °C for four days still showed a typical cold shock response with no significant difference in the percentage of cells exhibiting a response (Fig. 6D-E).

Taken together, these experiments do not indicate a direct requirement for the [Ca^2+^]_cyt_ elevations in protection from rapid cooling alone, as inhibition of the signalling response did not adversely affect physiology or growth following a cold shock, and the signalling response was not altered in cells acclimated to low temperatures.

### Interaction between cold and hypo-osmotic shock Ca^2+^ signalling pathways

Diatoms inhabiting inter-tidal regions may regularly experience a cold shock during tidal cycles (Fig 1), but this is unlikely to represent an isolated stressor. In particular, warming of shallow tidal pools can greatly increase their salinity due to evaporation (Firth and Williams, 2009), leading to a significant hypo-osmotic shock when the incoming tide reaches the tidal pool. *P. tricornutum* is highly perceptive to hypo-osmotic shock, exhibiting a large transient [Ca^2+^]_cyt_ elevation similar to those induced by cold shock (Falciatore et al., 2000; Helliwell et al., 2021). Since cold and hypo-osmotic shocks are likely to regularly co-occur in intertidal environments, we examined cellular Ca^2+^ signalling when these stressors were applied simultaneously.

A relatively mild hypo-osmotic shock (100% ASW to 95% ASW) administered to cells at 25 °C resulted in a single [Ca^2+^]_cyt_ elevation, as observed previously (Helliwell et al., 2021) (Fig. 7A). When the same hypo-osmotic shock was applied simultaneously with a cold shock (25 °C to 10 °C), both the amplitude and duration of the [Ca^2+^]_cyt_ elevations was substantially increased, although the number of cells exhibiting [Ca^2+^]_cyt_ elevations did not change (Fig. 7A-C). Hypo-osmotic shocks cause an increase in cell volume in *P. tricornutum*, which likely initiates [Ca^2+^]_cyt_ elevations through the activation of mechanosensitive ion channels (Helliwell et al., 2021). However, cell volume did not increase during cold shock (Supplemental Fig. S3), indicating that the rapid cooling does not simply elicit [Ca^2+^]_cyt_ elevations by mimicking a hypo-osmotic stimulus.

**Figure 7:**
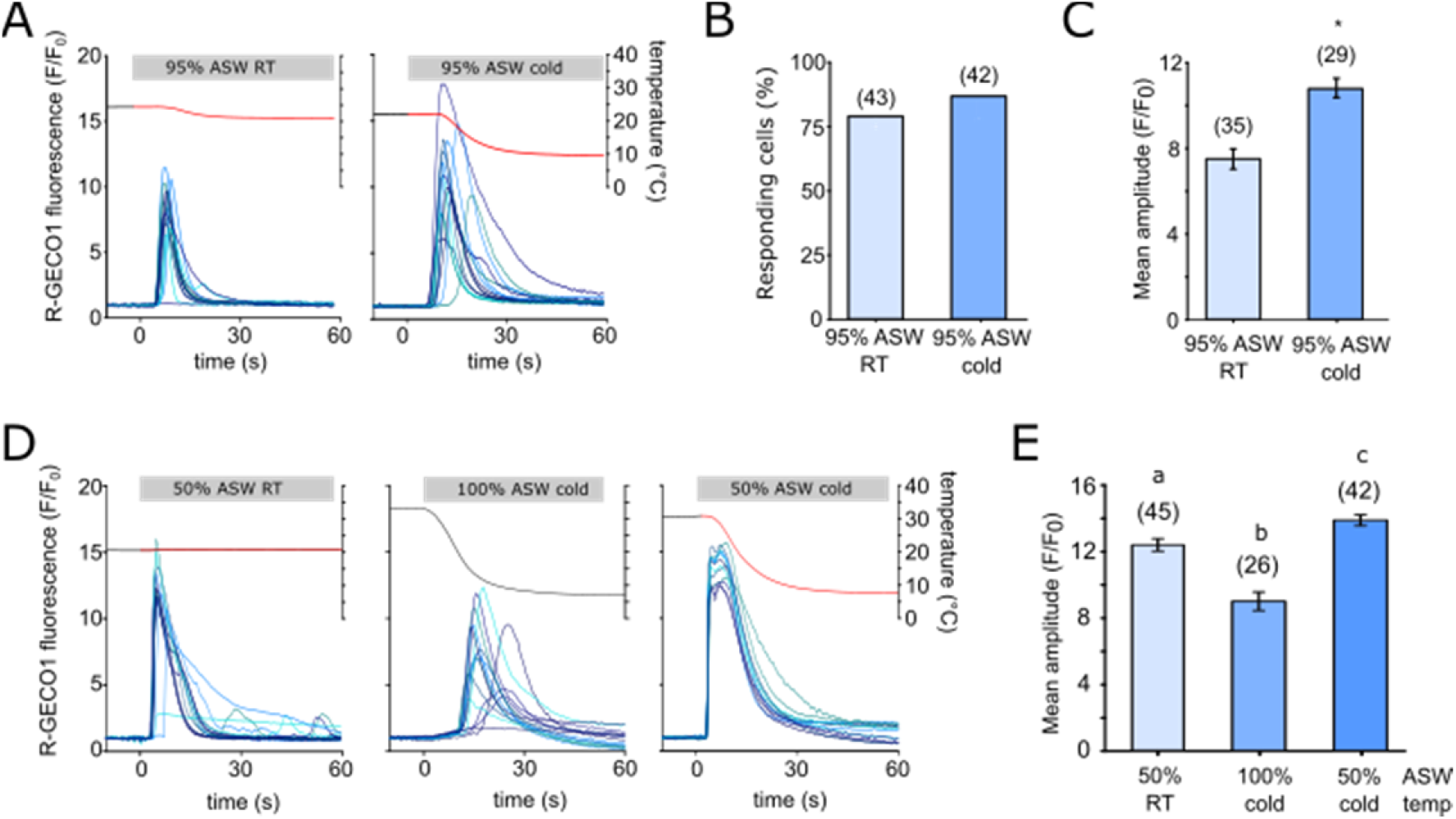
Interactions between the cold shock and hypo-osmotic shock Ca^2+^signalling pathways. **A)** R-GECO1 fluorescence ratio (F/F_0_) of PtR1 cells in response to a mild hypo-osmotic shock (95% ASW, left panel) or a simultaneous hypo-osmotic and cold shock (10 °C decrease, right panel). 12 representative traces are shown. **B)** Percentage of cells exhibiting [Ca^2+^]_cyt_ elevations for the experiment described in (A). Data are compiled from a minimum of two independent treatments. **C)** Mean amplitude of [Ca^2+^]_cyt_ elevations from responding cells in (B). The two treatments are significantly different (Student’s t-test P < 0.001). **D)** R-GECO1 fluorescence ratio of PtR1 cells in response to stronger simultaneous cold- and hypo-osmotic shocks. Cells were treated with a single hypo-osmotic shock (50% ASW), a single cold shock (10 °C) or a simultaneous cold- and hypo-osmotic shock (50% ASW, 10 °C). 13 representative traces are shown. **E)** Mean maximal amplitude of cells exhibiting [Ca^2+^]_cyt_ elevations in (D). For biphasic peaks the higher amplitude was chosen. The data represent the combination of at least three independent experiments per treatment. Letters represent significant differences between treatments (1-way Kruskal-Wallis ANOVA Ranks P < 0.001, with Dunn post hoc).

A stronger hypo-osmotic shock (100% ASW to 50 % ASW) resulted in a rapid [Ca^2+^]_cyt_ elevation which initiated directly after the stimulus was applied (Fig. 7D). In comparison, application of a cold shock from 34 °C to 8 °C triggered [Ca^2+^]_cyt_ elevations that rose less rapidly and exhibited a longer delay to their initiation (Fig. 7D). Combining both shocks using perfusion with 50% ASW at 10 °C led to biphasic [Ca^2+^]_cyt_ elevations in 71 % of cells (42 cells, three separate experiments) (Fig. 7D). These consisted of a very rapid initial peak in [Ca^2+^]_cyt_, followed by a second peak around 3 s later, which was of greater amplitude than the first peak in the majority of cells (24 out of 30). The mean maximal amplitude of the [Ca^2+^]_cyt_ elevations caused by the three different treatments were all significantly different from each other, with the cold shock alone causing the lowest and the combined cold- and hypo-osmotic shock causing the highest [Ca^2+^]_cyt_ elevations (Fig. 7E).

Taken together, [Ca^2+^]_cyt_ elevations induced by hypo-osmotic shock exhibit significant differences in amplitude and timing in the presence of a simultaneous cold shock. This indicates that the cold shock stimulus is additive and of sufficient magnitude to influence cellular Ca^2+^ signalling during hypo-osmotic stress. We therefore investigated whether Ca^2+^ signalling during cold shock may influence the short -term survival of *P. tricornutum* under hypo-osmotic stress.

### Simultaneous cold shock enhances survival during hypo-osmotic shock

Cells were treated with 25% ASW to administer a strong hypo-osmotic shock at control and low temperatures in the presence or absence of external Ca^2+^. Cell viability was determined after 3 min by staining with Sytox-Green. Administration of a cold shock alone, either in the presence or absence of external Ca^2+^, did not reduce cell viability (Fig 8). Application of a strong hypo-osmotic shock (25% ASW) significantly reduced cell viability, and this effect was greater following the removal of external Ca^2+^, supporting our previous observations that Ca^2+^ signalling is required for osmoregulation and volume control in *P. tricornutum* (Helliwell et al., 2021). Surprisingly, application of 25% ASW in combination with a cold shock (4 °C) led to a substantial reduction in cell mortality caused by hypo-osmotic shock (compared to the control temperature, 21°C). This effect was reduced by inhibiting Ca^2+^ signalling, although cell viability remained higher than at control temperature. Our data therefore indicate that rapid cooling has an important beneficial influence on the survival of *P. tricornutum* cells during a hypo-osmotic shock.

**Figure 8:**
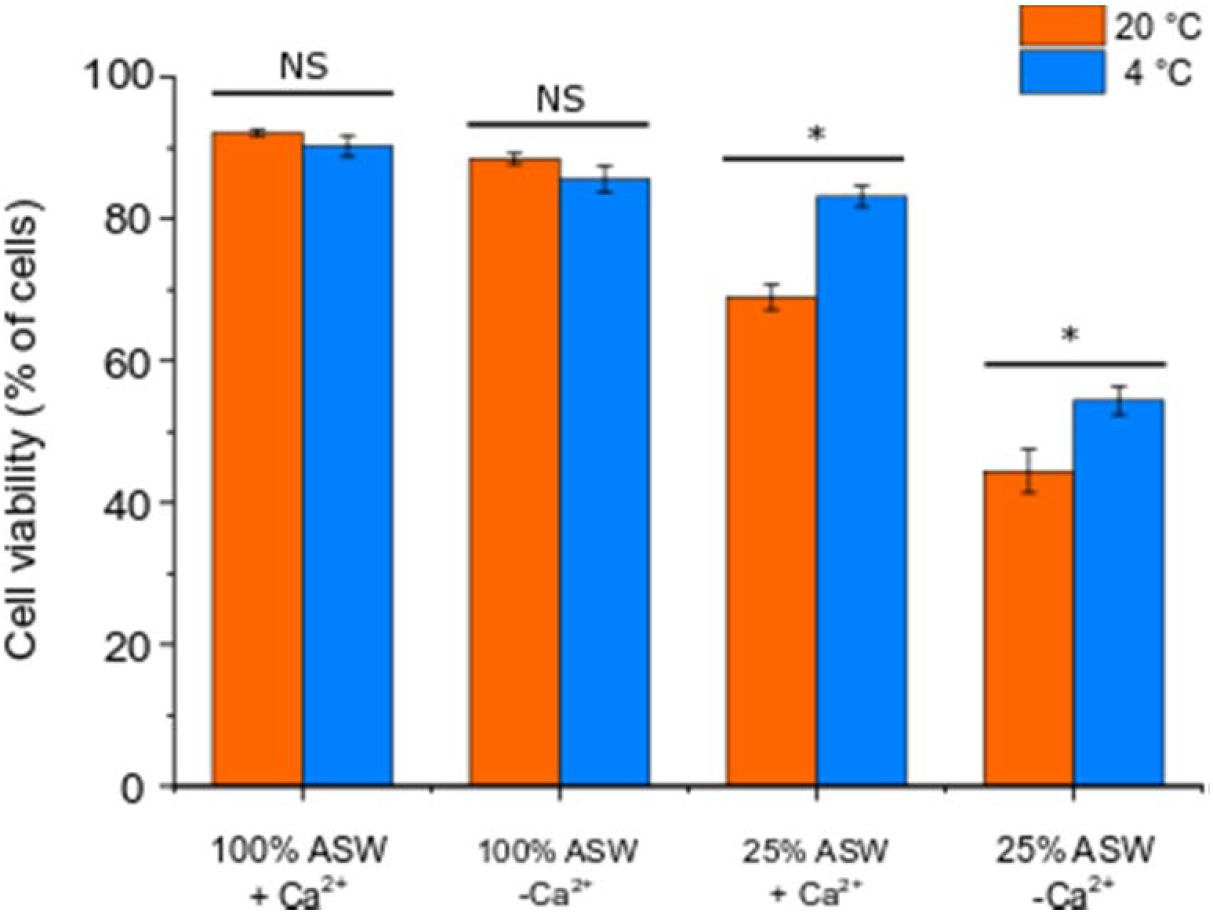
Simultaneous cold shock reduces mortality associated with hypo-osmotic shock. Cell viability (measured by exclusion of Sytox Green stain) was determined in *P. tricornutum* cells 3 min after exposure to a severe hypo-osmotic shock (25% ASW) with or without simultaneous cold shock (4 °C ASW). The presence or absence of external Ca^2+^ was used to establish the effect of inhibiting Ca^2+^ signalling during the applied shocks. The decrease in cell viability due to a hypo-osmotic shock (25% ASW) is significantly reduced when a simultaneous cold shock is applied. Three replicates were performed for each treatment, with at least 100 cells counted for each replicate. Significant differences due to temperature are marked with * (P<0.05 1-way ANOVA with Holm-Sidak post-hoc test). The experiment was repeated four times with similar results each time. Error bars represent SE.

### Cold shock is associated with Ca^2+^-dependent K^+^ efflux

Ca^2+^-dependent K^+^ efflux plays an essential role in cellular volume control in *P. tricornutum* during hypo-osmotic shock (Helliwell et al., 2021). We therefore tested whether the [Ca^2+^]_cyt_ elevations induced by cold shock also resulted in a K^+^ efflux that could influence cellular osmolarity. We settled a mono-layer of *P. tricornutum* cells onto a microscopy dish and used a K^+^-selective microelectrode to measure changes in extracellular K^+^ in the immediate vicinity of these cells. The cells were perfused with ASW at 25 °C, before rapidly switching to 12 °C. In each case, the cold shock induced a clear increase in extracellular K^+^ around the *P. tricornutum* cells (Fig 9A-C). Application of a cold shock in the absence of external Ca^2+^ greatly reduced K^+^ efflux from the cells, indicating that the K^+^ efflux is Ca^2+^-dependent. Very little change in extracellular K^+^ was observed during a cold shock in the absence of cells, indicating that the performance of the K^+^-selective microelectrode was not affected by the change in temperature. We conclude that cold shock induces Ca^2+^-dependent K^+^ efflux in *P. tricornutum* cells, which may contribute to volume regulation during a simultaneous hypo-osmotic shock.

**Figure 9:**
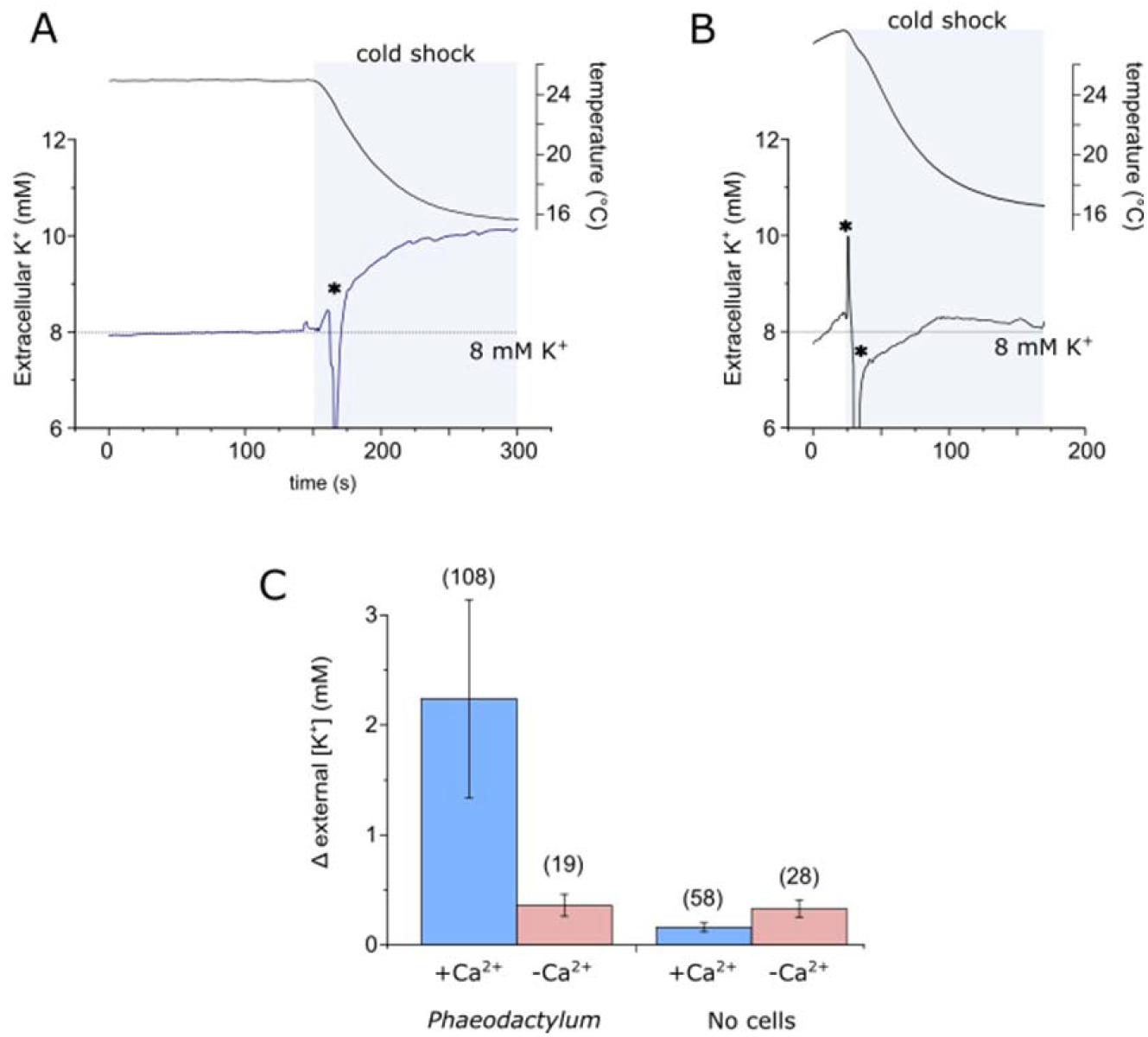
Cold shock induces a Ca^2+^-dependent K^+^ efflux. **A)** K^+^ efflux from *P. tricornutum* cells during a cold shock. A K^+^ microelectrode was placed adjacent to densely-packed *P. tricornutum* cells to measure K+ in the immediate vicinity of the cell. A cold shock was applied by perfusion. The increase in extracellular K^+^ is the result of K^+^ efflux from the cells. The temperature in the dish is also shown (upper trace). **B)** Extracellular K^+^ during a cold shock in the absence of external Ca^2+^ (perfusion with ASW-Ca^2+^ + 200 μM EGTA). **C)** Mean change in extracellular K^+^ around *P. tricornutum* cells during a cold shock. White bars indicate control experiments where the experimental set up was identical, but no *P. tricornutum* cells were present in order to assess whether the performance of the K^+^ microelectrode was influenced by temperature. The total number of replicates for each treatment are shown in parentheses, error bars = SE.

## Discussion

This study shows that physiologically significant transient [Ca^2+^]_cyt_ elevations are a consistent response to the rapid cold shocks likely to be experienced by intertidal diatoms. By using a continuous perfusion system, our study was able to avoid a shear-related [Ca^2+^]_cyt_ response, which may have masked a cold [Ca^2+^]_cyt_ response in earlier investigations using *P. tricornutum* expressing aequorin (Falciatore et al., 2000). The cold-induced [Ca^2+^]_cyt_ elevations are shown to be specifically involved in sensing the rate of cooling rather than the absolute temperature. A similar dependence of the amplitude of [Ca^2+^]_cyt_ elevations on the rate of cooling has been observed in *Arabidopsis*, which showed [Ca^2+^]_cyt_ elevations at cooling rates down to 0.05 °C s^-1^, (Plieth et al., 1999) indicating greater sensitivity of *Arabidopsis* to slower cooling rates. *P. tricornutum* and *T. pseudonana* did not show a Ca^2+^ signalling response to rapid warming, suggesting that the Ca^2+^ signalling pathways of animals and plants in response to elevated temperatures are not conserved in diatoms. Diatoms therefore likely use alternative cellular mechanisms for thermosensation in response to rapid heat shock, although as only short-term temperature increases were evaluated in our study, we cannot rule out a potential role for Ca^2+^ signalling in response to longer term temperature increases.

Cold shock-induced [Ca^2+^]_cyt_ elevations in *P. tricornutum* do not play an obvious role in acclimation to low temperatures. We found no longer-term growth effects of experimentally blocking the cold shock Ca^2+^ signal. Cold signalling in *P. tricornutum* therefore differs from plants and insects (Knight and Knight, 2012; Teets et al., 2013), in which the [Ca^2+^]_cyt_ elevations play a direct role in acclimation to lower temperatures. The [Ca^2+^]_cyt_ response in *P. tricornutum* is specifically induced by rapid cooling, which points to a potential role in short-term regulation of cellular processes rather than longer term acclimation to a change in temperature. Of particular interest is the interaction between cold shock and osmotic shock, since inter-tidal organisms are often likely to experience these stresses simultaneously, during an incoming tide or rain precipitation (Lewin and Guillard, 1963; Kirst, 1990). Given the nature of the osmotic and cold shock Ca^2+^ signals identified in *P. tricornutum*, it is most likely that they involve distinct sensory pathways, as evidenced by their additive nature and the appearance of a biphasic [Ca^2+^]_cyt_ elevation when cells were treated with simultaneous cold and osmotic shocks. Whether these distinct responses represent Ca^2+^ entry through different Ca^2+^ channels or are due to sequential activation of the same Ca^2+^ channel by different stimuli with little or no refractory period remains to be determined, although it is worthy of note that both the osmotic (Helliwell et al., 2021) and the cold-induced Ca^2+^ signals initiate at the cell apices (Fig 2B). Cold and osmotic Ca^2+^ signals also both require the presence of external Ca^2+^, indicating a shared requirement for plasma membrane Ca^2+^ channels, at least in the initiation of the [Ca^2+^]_cyt_ elevation.

The protective effect of cold shock on survival of *P. tricornutum* in response to severe osmotic shock may arise directly from cooperative Ca^2+^ signalling (Supplemental Fig. S4). The hypo-osmotic shock induced [Ca^2+^]_cyt_ elevations lead to rapid efflux of K^+^ in *P. tricornutum*, which restricts cell volume increase and prevents bursting (Helliwell et al., 2021). The results here strongly suggest that cold-induced [Ca^2+^]_cyt_ elevations may also act directly to trigger K^+^ efflux from the cytosol, for example through the activation of Ca^2+^-dependent K^+^ channels. Whether the rapid loss of K^+^ plays a physiological role in acclimation to low temperature is unclear, but it would clearly serve to lower the osmolarity of the cell. Given the frequent co-occurrence of cold and hypo-osmotic shocks, the cold-induced [Ca^2+^]_cyt_ elevations may therefore function primarily to support osmoregulation. Rapid cooling does not appear to adversely harm the cell when Ca^2+^ signalling is inhibited, whereas a severe hypo-osmotic shock will lead to cell bursting within seconds if cell volume is not controlled (Helliwell et al., 2021).

Osmoregulation in response to hypo-osmotic stress in diatoms (and most other eukaryotes) is most likely initiated by activation of mechanosensitive channels due to the increase in cell volume (Helliwell et al., 2021). Mechanosensitive channels only activate when the membrane is under tension, i.e. when swelling has already occurred, and cell viability is therefore under immediate threat if rapid osmoregulation cannot be achieved. The K^+^ efflux in response to a cold shock would allow the cell to reduce its osmolarity even if this critical increase in membrane tension is not perceived. By associating K^+^ efflux with an additional stimulus that commonly co-occurs with hypo-osmotic shock, diatoms can augment the osmoregulatory response and help prevent cell swelling to critical levels. Consistent with this hypothesis, cold-induced [Ca^2+^]_cyt_ elevations were only associated with very rapid cooling. A more gradual exposure to hypo-osmotic stress conveys a much lower risk of cell bursting, reducing the need to augment the osmoregulatory response.

We should also consider that low temperature may have a direct effect on reducing mortality during hypo-osmotic stress that is independent of the signalling component, for example by increasing cell wall rigidity. However, the protective effect of cold shock in the absence of Ca^2+^ was small compared to the much greater reduction in mortality in the presence of external Ca^2+^. We were unable to identify pharmacological inhibitors to selectively inhibit either osmotic or cold associated Ca^2+^ signalling and the removal of external Ca^2+^ completely inhibited both signalling pathways. Dissecting the individual contributions of these signalling pathways to cell survival during simultaneous shocks is therefore not currently easily achieved. Selective inactivation of the underlying molecular mechanisms through genetic approaches will most likely be required to fully understand the nature of the cross-talk between the signalling pathways.

Cellular responses to stressors are commonly examined in isolation in the laboratory in order to simplify the elucidation of the signalling pathways responsible. However, organisms often have to respond to inputs from multiple stimuli simultaneously in their natural environment, leading to cross-talk between signalling pathways. Cross-talk in cell signalling can occur when two distinct stimuli trigger a shared cellular response that confers tolerance to both stressors. This may involve activation of a common receptor or activation of independent receptors that converge on a specific node in the signalling pathway (Knight and Knight, 2001). Cross-talk with temperature sensing is likely to have evolved when another stress occurs simultaneously with temperature or with a predictable temporal link (i.e. one stimulus consistently precedes the other) (Sinclair et al., 2013). In the case of the inter-tidal zone, many environmental parameters will exhibit a degree of co-variance associated with tidal immersion and emersion. It seems likely that organisms inhabiting these environments have developed mechanisms of cross-talk in their pathways of environmental perception that enable them optimise their physiological responses.

There are multiple examples of cross-talk between temperature and osmotic stress signalling pathways in other eukaryotes. In plants, freezing temperatures can lead to cellular water loss due to external ice formation and many of the genes within the cold–inducible COR regulon are also inducible by drought (Boyce et al., 2003). The cold-responsive CBF/DREB1 and drought-responsive DREB2 transcription factors both bind to a common promoter element (DRE), leading to convergence of the cold and drought signalling pathways (Boyce et al., 2003). Overexpression of the cold-responsive DREB1A transcription factor in *Arabidopsis* resulted in enhanced tolerance to both freezing and drought stress (Liu et al., 1998). In addition, *Arabidopsis* plants treated with the phytohormone abscisic acid, which plays a primary role in drought tolerance, also show enhanced freezing tolerance (Mantyla et al., 1995). Cross-talk between temperature and osmotic stress signalling pathways have also been documented in yeast. *Saccharomyces cerevisiae* exhibits a high osmolarity (HOG) response to hyper-osmotic stress that results in increased production of the compatible solute, glycerol.

The HOG response is mediated by a mitogen-activated protein kinase (MAPK) pathway that can also be activated by other stimuli including both cold and heat shocks. Heat shock activates the HOG pathway indirectly by stimulating loss of glycerol, leading to hyper-osmotic stress (Winkler et al., 2002; Dunayevich et al., 2018).

Our results indicate that cross-talk between Ca^2+^-mediated cellular signalling mechanisms is an important consideration in the response of marine organisms to multiple stressors. Whilst our results are discussed primarily in the context of the inter-tidal zone where rapid substantial changes in temperature are a regular occurrence, the conserved nature of cold-induced Ca^2+^ signalling in *T. pseudonana* suggests that this pathway may be important more widely in diatom ecology. The cold-induced [Ca^2+^]_cyt_ elevations in *T. pseudonana* exhibit different characteristics from *P. tricornutum* that may reflect differences in their physiological response. Planktonic diatoms will undoubtedly encounter significant fluctuations in temperature and salinity in near-shore and estuarine environments or when they are mixed through the thermocline, although the magnitude and rate of the temperature changes are likely to be lower. Diatoms inhabiting sea ice environments may also experience rapid changes in temperature and salinity e.g. during flushing of hyper-saline brine channels with melt water (Mock and Junge, 2007). Future elucidation of the mechanisms of cross talk in these signalling pathways will allow us to understand how diatoms successfully integrate inputs from multiple environmental stimuli, which has likely played a major role in their success in diverse and highly dynamic environmental regimes.

## Acknowledgements

The work was supported by an ERC Advanced Grant to CB (ERC-ADG-670390) and a NERC award to GLW (NE/V013343/1).

## Author contributions

CB, FK and GLW conceived the study. FK performed the majority of the experimental analyses, including all imaging experiments. KEH contributed to measurements of osmotic stress. AC performed K^+^ microelectrode measurements. HPW and NM performed the environmental monitoring. TM, AH and TG contributed to the transformation of *T. pseudonana* with R-GECO1. GLW, CB and FK analysed the data and wrote the manuscript.

## Parsed Citations

Alverson AJ, Beszteri B, Julius ML, Theriot EC (2011) The model marine diatom Thalassiosira pseudonana likely descended from a freshwater ancestor in the genus Cyclotella. BMC Evol Biol 11: 125

Anderson SI, Rynearson TA (2020) Variability approaching the thermal limits can drive diatom community dynamics. Limnology and Oceanography 65: 1961–1973

Andrade EL, Luiz AP, Ferreira J, Calixto JB (2008) Pronociceptive response elicited by TRPA1 receptor activation in mice. Neuroscience 152: 511–520

Arrigoni C, Rohaim A, Shaya D, Findeisen F, Stein RA, Nurva SR, Mishra S, McHaourab HS, Minor DL, Jr. (2016) Unfolding of a temperature-sensitive domain controls voltage-gated channel activation. Cell 164: 922–936

Baker KG, Geider RJ (2021) Phytoplankton mortality in a changing thermal seascape. Glob Chang Biol

Boyce JM, Knight H, Deyholos M, Openshaw MR, Galbraith DW, Warren G, Knight MR (2003) The sfr6 mutant of Arabidopsis is defective in transcriptional activation via CBF/DREB1 and DREB2 and shows sensitivity to osmotic stress. Plant J 34: 395–406

Christensen AP, Akyuz N, Corey DP (2016) The outer pore and selectivity filter of TRPA1. PLoS One 11: e0166167

Clapham DE, Miller C (2011) Athermodynamic framework for understanding temperature sensing by transient receptor potential (TRP) channels. Proc Natl Acad Sci U S A 108: 19492–19497

Cui Y, Lu S, Li Z, Cheng J, Hu P, Zhu T, Wang X, Jin M, Wang X, Li L, Huang S, Zou B, Hua J (2020) Cyclic nucleotide-gated ion channels 14 and 16 promote tolerance to heat and chilling in rice. Plant Physiol 183: 1794–1808

De Martino A, Bartual A, Willis A, Meichenin A, Villazan B, Maheswari U, Bowler C (2011) Physiological and molecular evidence that environmental changes elicit morphological interconversion in the model diatom Phaeodactylum tricornutum. Protist 162: 462–481

De Martino A, Meichenin A, Shi J, Pan KH, Bowler C (2007) Genetic and phenotypic characterization of Phaeodactylum tricornutum (Bacillariophyceae) accessions. Journal of Phycology 43: 992–1009

Dunayevich P, Baltanas R, Clemente JA, Couto A, Sapochnik D, Vasen G, Colman-Lerner A (2018) Heat-stress triggers MAPK crosstalk to turn on the hyperosmotic response pathway. Sci Rep 8: 15168

Falciatore A, d’Alcala MR, Croot P, Bowler C (2000) Perception of environmental signal by a marine diatom. Science 288: 2363–2366

Firth LB, Williams GA (2009) The influence of multiple environmental stressors on the limpet Cellana toreuma during the summer monsoon season in Hong Kong. Journal of Experimental Marine Biology and Ecology 375: 70–75

Furuya T, Matsuoka D, Nanmori T (2014) Membrane rigidification functions upstream of the MEKK1-MKK2-MPK4 cascade during cold acclimation in Arabidopsis thaliana. FEBS Lett 588: 2025–2030

Gattuso JP, Magnan A, Bille R, Cheung WWL, Howes EL, Joos F, Allemand D, Bopp L, Cooley SR, Eakin CM, Hoegh-Guldberg O, Kelly RP, Portner HO, Rogers AD, Baxter JM, Laffoley D, Osborn D, Rankovic A, Rochette J, Sumaila UR, Treyer S, Turley C (2015) Contrasting futures for ocean and society from different anthropogenic CO2 emissions scenarios. Science 349

Gruber N, Boyd PW, Frolicher TL, Vogt M (2021) Biogeochemical extremes and compound events in the ocean. Nature 600: 395–407

Guillard RL (1975) Culture of phytoplankton for feeding marine invertebrates. Culture of Marine Invertebrate Animals: 29–60

Harley CDG, Hughes AR, Hultgren KM, Miner BG, Sorte CJB, Thornber CS, Rodriguez LF, Tomanek L, Williams SL (2006) The impacts of climate change in coastal marine systems. Ecology Letters 9: 228–241

Helliwell KE, Chrachri A, Koester JA, Wharam S, Verret F, Taylor AR, Wheeler GL, Brownlee C (2019) Alternative mechanisms for fast Na+/Ca2+ signaling in eukaryotes via a novel class of single-domain voltage-gated channels. Curr Biol 29: 1503–1511 e1506

Helliwell KE, Harrison EL, Christie-Oleza JA, Rees AP, Kleiner FH, Gaikwad T, Downe J, Aguilo-Ferretjans MM, Al-Moosawi L, Brownlee C, Wheeler GL (2021) ANovel Ca2+ signaling pathway coordinates environmental phosphorus sensing and nitrogen metabolism in marine diatoms. Curr Biol 31: 978–989 e974

Helliwell KE, Kleiner FH, Hardstaff H, Chrachri A, Gaikwad T, Salmon D, Smirnoff N, Wheeler GL, Brownlee C (2021) Spatiotemporal patterns of intracellular Ca2+ signalling govern hypo-osmotic stress resilience in marine diatoms. New Phytol 230: 155–170

J C Lewin a, Guillard RRL (1963) Diatoms. Annual Review of Microbiology 17: 373–414

Javaheri N, Dries R, Burson A, Stal LJ, Sloot PMA, Kaandorp JA (2015) Temperature affects the silicate morphology in a diatom. Scientific Reports 5: 11652 .

Kirst GO (1990) Salinity tolerance of eukaryotic marine-algae. Annual Review of Plant Physiology and Plant Molecular Biology 41: 21–53

Knight H, Knight MR (2001) Abiotic stress signalling pathways: specificity and cross-talk. Trends Plant Sci 6: 262–267

Knight H, Trewavas AJ, Knight MR (1996) Cold calcium signaling in Arabidopsis involves two cellular pools and a change in calcium signature after acclimation. Plant Cell 8: 489–503

Knight MR, Knight H (2012) Low-temperature perception leading to gene expression and cold tolerance in higher plants. New Phytol 195: 737–751

Lenzoni G, Knight MR (2019) Increases in absolute temperature stimulate free calcium concentration elevations in the chloroplast. Plant Cell Physiol 60: 538–548

Lewin JC, Guillard RRL (1963) Diatoms. Annual Review of Microbiology 17: 373–414

Liang Y, Koester JA, Liefer JD, Irwin AJ, Finkel ZV (2019) Molecular mechanisms of temperature acclimation and adaptation in marine diatoms. ISME J 13: 2415–2425

Liu Q, Ding Y, Shi Y, Ma L, Wang Y, Song C, Wilkins KA, Davies JM, Knight H, Knight MR, Gong Z, Guo Y, Yang S (2021) The calcium transporter ANNEXIN1 mediates cold-induced calcium signaling and freezing tolerance in plants. EMBO J 40: e104559

Liu Q, Kasuga M, Sakuma Y, Abe H, Miura S, Yamaguchi-Shinozaki K, Shinozaki K (1998) Two transcription factors, DREB1 and DREB2, with an EREBP/AP2 DNAbinding domain separate two cellular signal transduction pathways in drought-and low-temperature-responsive gene expression, respectively, in Arabidopsis. Plant Cell 10: 1391–1406

Ma Y, Dai X, Xu Y, Luo W, Zheng X, Zeng D, Pan Y, Lin X, Liu H, Zhang D, Xiao J, Guo X, Xu S, Niu Y, Jin J, Zhang H, Xu X, Li L, Wang W, Qian Q, Ge S, Chong K (2015) COLD1 confers chilling tolerance in rice. Cell 160: 1209–1221

Malviya S, Scalco E, Audic S, Vincent F, Veluchamy A, Poulain J, Wincker P, Iudicone D, de Vargas C, Bittner L, Zingone A, Bowler C (2016) Insights into global diatom distribution and diversity in the world’s ocean. Proc Natl Acad Sci U S A 113: E1516–1525

Mantyla E, Lang V, Palva ET (1995) Role of abscisic acid in drought-induced freezing tolerance, cold acclimation, and accumulation of LT178 and RAB18 proteins in Arabidopsis thaliana. Plant Physiol 107: 141–148

Mock T, Junge K (2007) PSYCHROPHILIC DIATOMS: Mechanisms for survival in freeze–thaw cycles. In J Seckbach, ed, Algae and Cyanobacteria in Extreme Environments. Springer, pp 343–364

Montagnes DJS, Franklin DJ (2001) Effect of temperature on diatom volume, growth rate, and carbon and nitrogen content: Reconsidering some paradigms. Limnology and Oceanography 46: 2008–2018

Ohkura M, Sasaki T, Sadakari J, Gengyo-Ando K, Kagawa-Nagamura Y, Kobayashi C, Ikegaya Y, Nakai J (2012) Genetically encoded green fluorescent Ca2+ indicators with improved detectability for neuronal Ca2+ signals. PLoS One 7: e51286

Plieth C, Hansen UP, Knight H, Knight MR (1999) Temperature sensing by plants: the primary characteristics of signal perception and calcium response. Plant J 18: 491–497

Pokorna J, Schwarzerova K, Zelenkova S, Petrasek J, Janotova I, Capkova V, Opatrny Z (2004) Sites of actin filament initiation and reorganization in cold-treated tobacco cells. Plant Cell and Environment 27: 641–653

Saidi Y, Finka A, Muriset M, Bromberg Z, Weiss YG, Maathuis FJ, Goloubinoff P (2009) The heat shock response in moss plants is regulated by specific calcium-permeable channels in the plasma membrane. Plant Cell 21: 2829–2843

Schaum CE, Buckling A, Smirnoff N, Studholme DJ, Yvon-Durocher G (2018) Environmental fluctuations accelerate molecular evolution of thermal tolerance in a marine diatom. Nat Commun 9: 1719

Sengupta P, Garrity P (2013) Sensing temperature. Curr Biol 23: R304–307

Silva DF, de Almeida MM, Chaves CG, Braz AL, Gomes MA, Pinho-da-Silva L, Pesquero JL, Andrade VA, Leite Mde F, de Albuquerque JG, Araujo IG, Nunes XP, Barbosa-Filho JM, Cruz Jdos S, Correia Nde A, de Medeiros IA (2015) TRPM8 channel activation induced by monoterpenoid rotundifolone underlies mesenteric artery relaxation. PLoS One 10: e0143171

Sinclair BJ, Ferguson LV, Salehipour-shirazi G, MacMillan HA (2013) Cross-tolerance and cross-talk in the cold: relating low temperatures to desiccation and immune stress in insects. Integr Comp Biol 53: 545–556

Smale DA, Wernberg T, Oliver ECJ, Thomsen M, Harvey BP, Straub SC, Burrows MT, Alexander LV, Benthuysen JA, Donat MG, Feng M, Hobday AJ, Holbrook NJ, Perkins-Kirkpatrick SE, Scannell HA, Sen Gupta A, Payne BL, Moore PJ (2019) Marine heatwaves threaten global biodiversity and the provision of ecosystem services. Nature Climate Change 9: 306–+

Souffreau C, Vanormelingen P, Verleyen E, Sabbe K, Vyverman W (2010) Tolerance of benthic diatoms from temperate aquatic and terrestrial habitats to experimental desiccation and temperature stress. Phycologia 49: 309–324

Svensson F, Norberg J, Snoeijs P (2014) Diatom cell size, coloniality and motility: trade-offs between temperature, salinity and nutrient supply with climate change. Plos One 9

Tahtiharju S, Sangwan V, Monroy AF, Dhindsa RS, Borg M (1997) The induction of kin genes in cold-acclimating Arabidopsis thaliana. Evidence of a role for calcium. Planta 203: 442–447

Teets NM, Yi SX, Lee RE, Jr., Denlinger DL (2013) Calcium signaling mediates cold sensing in insect tissues. Proc Natl Acad Sci U S A 110: 9154–9159

Vardi A, Formiggini F, Casotti R, De Martino A, Ribalet F, Miralto A, Bowler C (2006) Astress surveillance system based on calcium and nitric oxide in marine diatoms. Plos Biology 4: 411–419

Verret F, Wheeler G, Taylor AR, Farnham G, Brownlee C (2010) Calcium channels in photosynthetic eukaryotes: implications for evolution of calcium-based signalling. New Phytol 187: 23–43

Winkler A, Arkind C, Mattison CP, Burkholder A, Knoche K, Ota I (2002) Heat stress activates the yeast high-osmolarity glycerol mitogen-activated protein kinase pathway, and protein tyrosine phosphatases are essential under heat stress. Eukaryot Cell 1: 163–173

Xu H, Ramsey IS, Kotecha SA, Moran MM, Chong JA, Lawson D, Ge P, Lilly J, Silos-Santiago I, Xie Y, DiStefano PS, Curtis R, Clapham DE (2002) TRPV3 is a calcium-permeable temperature-sensitive cation channel. Nature 418: 181–186

Yin Y, Wu M, Zubcevic L, Borschel WF, Lander GC, Lee SY (2018) Structure of the cold-and menthol-sensing ion channel TRPM8. Science 359: 237–241

Zhao Y, Araki S, Wu J, Teramoto T, Chang YF, Nakano M, Abdelfattah AS, Fujiwara M, Ishihara T, Nagai T, Campbell RE (2011) An expanded palette of genetically encoded Ca2+ indicators. Science 333: 1888–1891

